# Metacommunity Structure Preserves Genome Diversity in the Presence of Gene-specific Selective Sweeps

**DOI:** 10.1101/2023.01.17.524383

**Authors:** Simone Pompei, Edoardo Bella, Joshua S. Weitz, Jacopo Grilli, Marco Cosentino Lagomarsino

**Affiliations:** IFOM ETS - The AIRC Institute of Molecular Oncology, Milan, Italy; Dipartimento di Fisica, Università degli Studi di Milano, and I.N.F.N, via Celoria 16 Milano, Italy; School of Biological Sciences, Georgia Institute of Technology, Atlanta, GA, USA; School of Physics, Georgia Institute of Technology, Atlanta, GA, USA; Institut de Biologie, École Normale Supérieure, Paris, France; Quantitative Life Sciences, The Abdus Salam International Centre for Theoretical Physics (ICTP), Trieste 34151, Italy

## Abstract

The horizontal transfer of genes is fundamental for the eco-evolutionary dynamics of microbial communities, such as oceanic plankton, soil, and the human microbiome. In the case of an acquired beneficial gene, classic population genetics would predict a genome-wide selective sweep, whereby the genome spreads clonally with the gene, removing genome diversity. Instead, several sources of metagenomic data show the existence of “gene-specific sweeps”, whereby a beneficial gene spreads across a bacterial community, maintaining genome diversity. Several hypotheses have been proposed to explain this process, including the decreasing gene flow between ecologically distant populations, frequency-dependent selection from linked deleterious allelles, and very high rates of horizontal gene transfer. Here, we propose an additional possible scenario grounded in eco-evolutionary principles. Specifically, we show by a mathematical model and simulations that a metacommunity where species can occupy multiple patches helps maintain genome diversity. Assuming a scenario of patches dominated by single species, our model predicts that diversity only decreases moderately upon the arrival of a new beneficial gene, and that losses in diversity can be quickly restored. We explore the generic behavior of diversity as a function of three key parameters, frequency of insertion of new beneficial genes, migration rates and horizontal transfer rates.Our results provides a testable explanation for how diversity can be maintained given gene-specific sweeps even in the absence of high horizontal gene transfer rates.

## INTRODUCTION

Horizontal Gene Transfer (HGT) plays a crucial role in the processes that shape bacterial evolution [1–5]. HGT accelerates the adaptation of bacterial communities to new ecological niches [6, 7] and reduces the deleterious effects associated with the accumulation of genetic load by clonal reproduction [8]. HGT is also a widespread pathway through which pathogenic bacteria acquire resistance to antibiotics [7, 9].

According to classical population genetics theories [10], when the rate of HGT is low, then vertical inheritance is the main mechanism for the expansion of novel genetic variants. In such case, the evolutionary dynamics is described by clonal evolution. In this case, because of linkage effects, transfer of a highly beneficial gene results in a drastic reduction of the diversity, as a consequence of the clonal expansion of the mutant carrying the gene, whose genome would “sweep” together with the beneficial gene. We can term this process a genome-wide selective sweep.

In contrast to this scenario, several lines of metagenomic evidence from communities of phylogenetically related strains [1, 11, 12], support an alternative scenario of gene-specific sweeps, an evolutionary dynamics where a beneficial gene can reach fixation across species or strains, without erasing diversity. Reconciling this scenario with standard population genetic models would require very high recombination rates [11]. Hence, the consensus is that more complex mechanisms should be in place [1].

In the past years, several alternative hypotheses have been put forward to reconcile the evidence of gene sweeps with the expected values of the recombination rate. A first mechanism, originally proposed by Shapiro and coworkers [11] uses the observation that HGT rates between pairs of species decline fast with their genetic distance [13]. This mechanism would lead to an effective HGT barrier between populations of different species/strains, provided that the intra-population rate of the genomic changes is faster than the inter-population HGT rate. Therefore diversity could persist in a metacommunity (a group of populations including different species/strains based on spatially separate patches) in presence of selective effects. A second mechanism, proposed by Niehus and coworkers [6], proposes that the combination of a small migration rate and a high HGT rate can lead to the fixation of beneficial genes without the decrease of genome diversity. A different hypothesis, put forward by Takeuchi and coworkers in 2015 [14], involves the linkage of the beneficial sweeping gene with widespread (species-specific) deleterious alleles, which would lead to negative frequency-dependent selection. This mechanism was shown to explain a gene-sweep dynamics in quantitative terms, provided that the basal recombination rate is sufficiently low. The widespread linkage of the beneficial gene with deleterious alleles does not have a simple explanation, but the authors speculate that it could be the consequence of ecological interactions between bacteria and viral predators [14], possibly supported by a “Kill the Winner” dynamics [15]. Finally, a complementary scenario for gene sweeping might be the so-called “soft sweeps” [16, 17], where widespread (e.g. neutral) recombination through HGT can generate a pool of standing variation (in our case promoting the presence of one or more beneficial alleles of a given gene across species) that is sufficient to support the emergence of multiple (interfering) sweeps in parallel upon a change of selective pressure. In this case the gene-specific sweep would consist of multiple parallel sweeps of a gene that previously spread neutrally by HGT. However, this scenario is based as the previous ones on high HGT rates, and cannot explain situations where a gene sweep originates from a gene that is not already initially present neutrally in many species. While the proposed mechanisms are very different, as a general rule, the assumption of high HGT rates appears to be crucial to most of them.

Here, we propose a complementary eco-evolutionary mechanism, given a metacommunity structure, which shows that high HGT is not required for gene sweeps. Specifically, we assume that the environment is characterized by the presence of multiple patches (e.g., nutrient patches as in marine snow [18]) and that their physical separation is the main limitation to the spread, through genome migration or HGT, of beneficial genes. As we show metacommunity structure can preserve diversity during the fixation dynamics of a beneficial gene without requiring high recombination rates.

## RESULTS

### Model components

We introduce an evolutionary model that describes the dynamics of the diversity in a spatially structured environment, supporting different species in the presence of gene sweeps. Our model describes a metacommunity where each species dominates in a single habitat (or in other words intra-habitat dynamics is assumed to lead to neutral or selective sweeps of a single species). In the following, we aim to describe a scenario where the intra-species genetic diversity is very limited (and restricted to neutral diversity only), so that all the individuals belonging to the same species can effectively be associated to a single strain (the consensus genome of the individuals of the same specie). Fig. 1 illustrates the key model ingredients. We consider M distinct patches, each of which supports a single phenotypically homogeneous population, i.e., it only contains a single species. The same species (corresponding, depending on the specific context, to ecotypes or strains) can populate more than a single patch. The diversity is quantified by the number *S* (1 ≤ *S* ≤ *M*) of distinct species present in the community at a given time-point.

**Figure 1.**
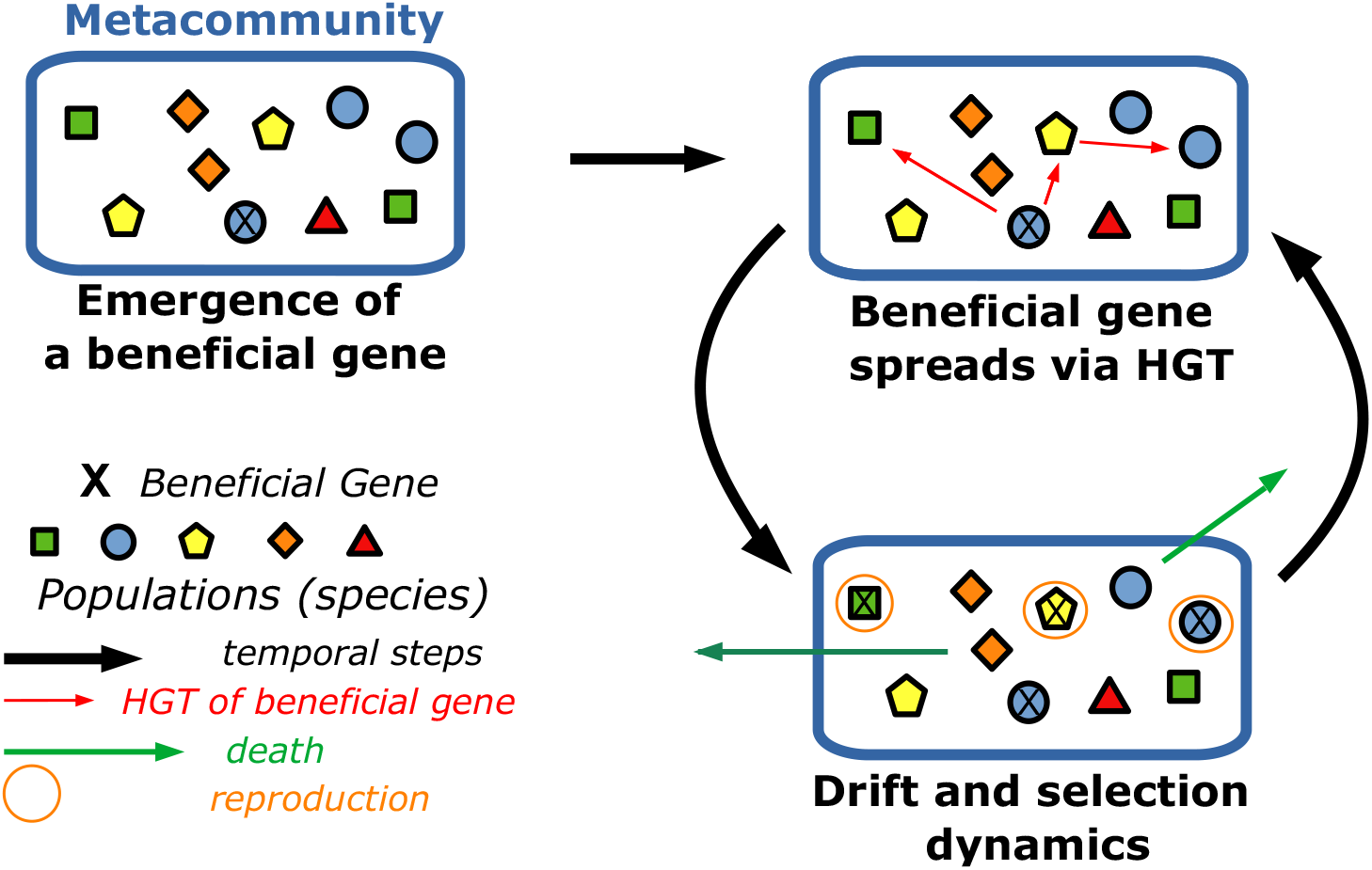
Schematic representation of the multi-species metacommunity and the temporal dynamics of the model. We consider a metacommunity consisting of *M* distinct patches (represented by symbols) and supporting different species (represented by colors/shapes). We assume that each patch is populated by only one species (a colored geometric shape in the picture). Species in the metacommunity can contain a beneficial gene in their genome, here schematically represented by a cross. The time evolution of the metacommunity is the result of a migration-selection dynamics across patches, and selective effects are associated to the presence-absence of beneficial genes. The spread of beneficial genes in the metacommunity is the result of HGT events between patches.

A beneficial gene can spread across patches by two mechanisms, migration of an individual (and its genome), with the consequent genome-wide sweep of the strains carrying the gene, or HGT and sweep of the gene in the community. In presence of a diversity-maintenance mechanism, enforced with a certain time scale, the diversity loss from genome-wide sweeps competes with the emergence dynamics of new species, inducing an increase of the diversity. In the model, as in standard Gillespie algorithmic approaches [19], time is counted in terms of the number of steps, and at the end of each step a single move will occur (e.g, a migration or an invasion). Hence, although each move is associated a physical rate, because of this normalization, these physical rates can be treated as probabilities (i.e., are dimensionless). Time is also counted in terms of an another unit, which we will be referred to as a “generation” of the metacommunity, where 1gen = *M* time steps.

### Dynamics of the diversity of the metacommunity in absence of HGT

In order to provide a mechanism for diversity-maintenance, we used a neutral process [20– 23] where the species occupation of patches change over time because of (i) neutral migration-sweep events, where a species occupying a patch is replaced by another existing species (ii) innovation events consisting in the emergence of a new species (which accounts for migration-sweep events from external species and speciation events).

Beneficial genes can be generated and spread across patches via HGT. We note that the specifics of the diversity-maintenance mechanism are not relevant for our results, which focus on the diversity loss (and time scale) due to the HGT-migration process of the beneficial gene. The only role of the diversity-maintenance process in our model is to provide a time scale that competes with the diversity loss due to gene sweeps. In the following, we will explore three distinct regimes of the dynamics of the model, which correspond to different combinations of the competing time scales. Our main result is that the diversity loss in presence of a beneficial gene can be moderate thanks to the presence of many patches supporting different species. Values of the rates associated to the different time scales will be defined and discussed in each regime.

We first focus on the evolutionary dynamics of the diversity-maintenance process, in the absence of HGT (Fig. 2A). This neutral model contains two elementary moves: (i) neutral migration/sweep across patches and (ii) innovation events, corresponding in this context ti the emergence of new species. At each time step, corresponding to the characteristic time scale of migration-sweep events, each population occupying a patch in the metacommunity can either change into a population corresponding to a species as a result of an innovation event (speciation or migration and neutral sweep from an outside pool of species), with a rate *ν* (per patch/per time step) or migrate and sweep neutrally to another patch, with a rate 1 − *ν*, (per patch/per time step) causing the extinction of the pre-existing species in the invaded patch (Fig. 2A).

**Figure 2.**
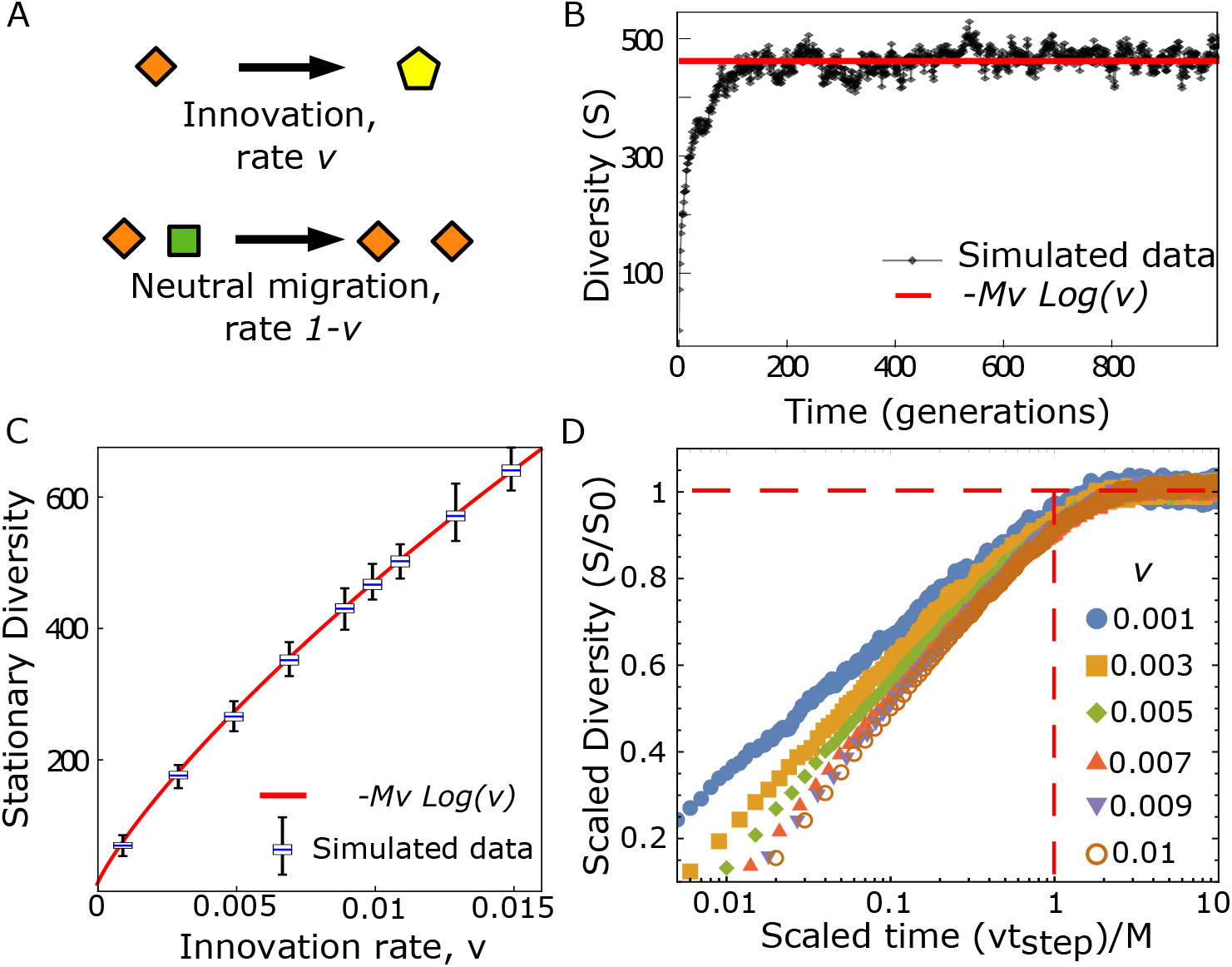
In absence of HGT, an infinite-allele model provides a diversity-maintenance mechanism with an intrinsic time scale. **A**. We consider a standard neutral model maintaining diversity. At each time step, corresponding to a basic migration-sweep time scale each patch can be swept neutrally by a new species, with an innovation event taking place with rate *ν* (per patch, per time step), or alternatively, its species can migrate and sweep, invading another patch (and sweeping neutrally), with a rate 1−*ν*. Panel **B** shows diversity (*S*(*t*), defined as the total number of distinct species present in the metacommunity at a given time *t*) in a typical simulation. Diversity displays an relaxation dynamics to a plateau, in agreement with (Eq. 1). The parameters of the simulation are *ν* = 0.01 and *M* = 10 000. **C**. Comparison between Eq. 1 (valid in the limit *M* ≫ 1 and *ν* ≪ 1) and simulated data for the equilibrium value of the diversity, plotted as a function of the innovation rate (*ν*). Simulations correspond to *M* = 10 000 and averages over 100 independent realizations. For each simulated value of *ν* the distribution of diversity *S* is shown as a box plot (blue line: mean value, box: inter-quartile range, fences: max and min values) **D**. Characterization of the relaxation time scale for the average trajectories of the diversity. The plot shows diversity (scaled by its equilibrium value) vs time, scaled by the common relaxation time scale *N/ν*. Simulations were performed over 100 independent realizations, for different values of the innovation rate *ν* (shown in the legend, coded by color and symbol). All the simulations were initialized with a metacommunity of a single species (*S*(*t* = 0) = 1).

In this regime, the diversity *S* displays a “relaxation to equilibrium” dynamics (Fig 2B), reaching a stationary state *S*_0_. The analytical expressions for the equilibrium value can be derived from classic calculations (see Methods), and is

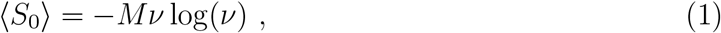

computed in the limit *M* ≫ 1 and *ν* ≪ 1. Our numerical simulations of the model (Fig. 2C) agree with Eq. 1, showing that the typical relaxation time to the equilibrium, for this model setting is the inverse of the innovation rate *τ*_*eq*_ ≃ *M/*(*ν*) steps = 1*/ν* gen (Fig 2D).

### Metacommunity diversity loss resulting from the introduction of a beneficial gene

We next ask when diversity decreases only moderately upon the introduction of a beneficial gene, even in the absence of diversity-maintenance mechanisms. In order to address this question, we focused on the dynamics following the introduction of a beneficial gene that can spread through the metacommunity via both HGT and sweep (gene-specific sweep) and genome-wide sweeps on single patches. The initial diversity was set using the neutral model described in the previous section (hence depends on *ν*, see Eq. 1). We considered the limiting situation where no diversity-restoring mechanism was in action during the whole sweep. In other words, we assume that the fixation dynamics of the advantageous gene in the metacommunity is much faster than the relaxation time scale of the neutral biodiversity model. Roughly, if there are *N* individuals per patch, this limit corresponds to the condition

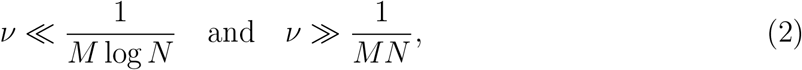

i.e., where the genome sweep time *M* log(*N*) is much smaller than 1*/ν*, but this is in turn much smaller than a metacommunity-wide fixation time, which is order *MN*. This assumption, which will be relaxed in the next paragraph, is the most conservative scenario (the most adverse in terms of diversity loss) for the introduction of a new beneficial gene in a metacommunity.

Under these assumptions, only two processes take place (Fig. 3A): (i) migration-sweep of a patch by a species carrying the beneficial gene (genome-wide sweep), which leads to a reduction of diversity, and (ii) spread of the beneficial gene by HGT (gene sweep), which does not reduce diversity. Hence, diversity can only decrease in this scenario, we are interested in the magnitude of the decrease relative to the diversity baseline (see details in the Methods, Eq. 7).

**Figure 3.**
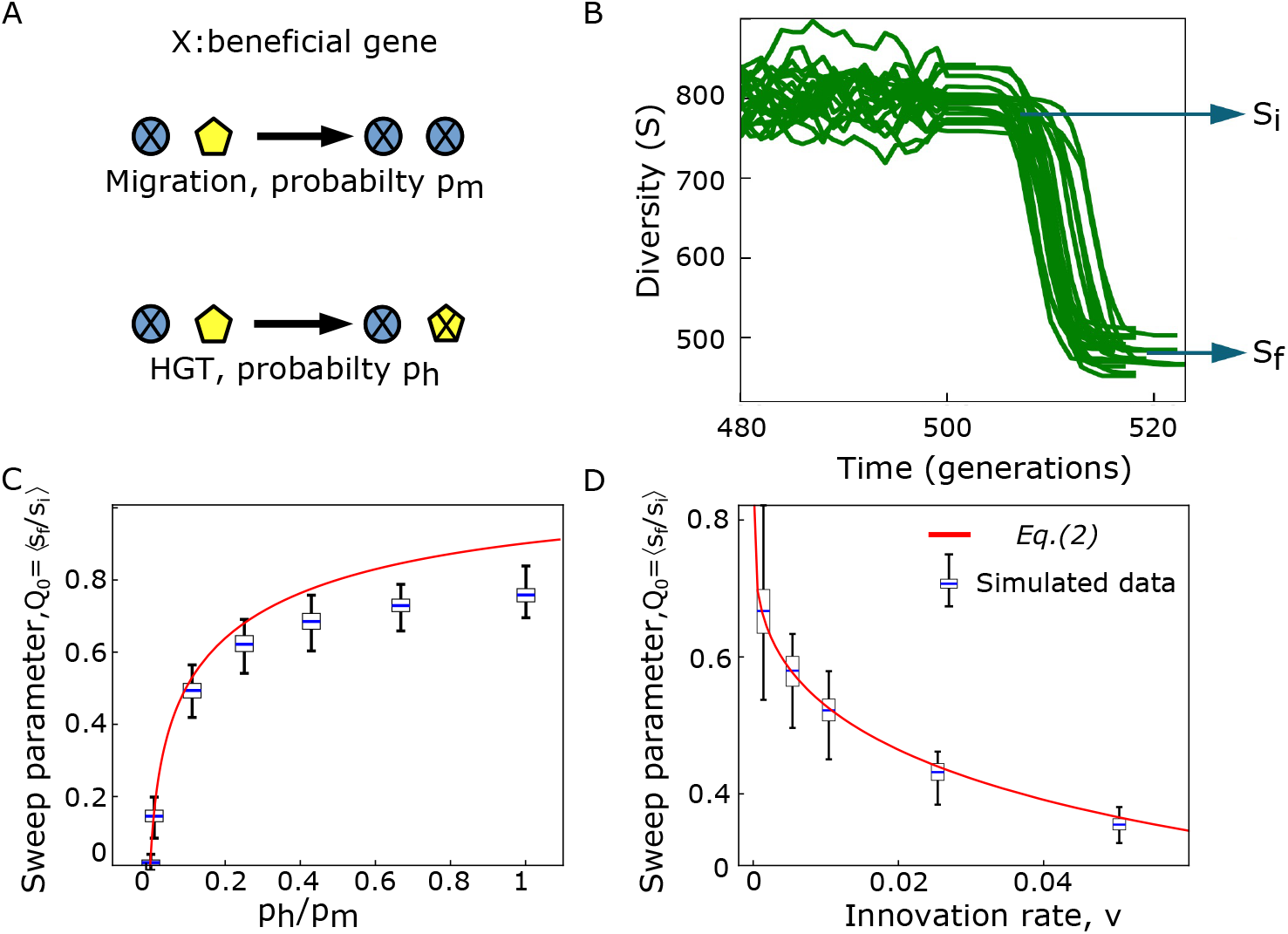
In presence of HGT only, and no diversity-maintenance mechanism, the fixation of a beneficial gene in the metacommunity can lead to a moderate loss of diversity. **A**. A beneficial gene can spread across patches (i) via migration events (reducing diversity) or (ii) via HGT-sweep (maintaining diversity) The two processes take place at each time step, with rate *p*_*m*_ (per patch, per time step) and *p*_*h*_ (per patch, per time step) respectively. Here, we assume that no diversity-maintenance mechanism counteracts the diversity loss (this assumption will be relaxed later) **B**. Simulations of this model show a diversity (*S*(*t*)) loss from the initial value (*S*_*i*_) to a new stable value (*S*_*f*_), corresponding to complete invasion of the beneficial gene. Each solid line is a realization, with initial condition of the simulations generated by the neutral model described in Fig. 2 (*M* = 10000 and *ν* = 0.02), and with parameters *p*_*h*_ = 0.2 and *p*_*m*_ = 1 − *p*_*h*_. **C-D**. Comparison between analytical prediction (Eq.3) and simulated data for the sweep parameter *Q* = ⟨*S*_*f*_ */S*_*i*_⟩ as a function of the ratio *p*_*h*_*/p*_*m*_ (panel **C**, *ν* = 0.01), and of the innovation rate of the neutral model generating the initial diversity (panel **D**, *p*_*h*_ = 0.1, *p*_*m*_ = 0.9). Simulations performed with *M* = 10 000 realizations. The panels **BC** show the distribution of diversity *S* over 100 realizations as a box plot (blue line: mean value, box: inter-quartile range, fences: max and min values).

To implement genome-wide migration-sweep events, at each time step, with a rate *p*_*m*_ (per patch, per time step), two patches are picked randomly. If the first patch carries the beneficial gene and the second one does not, the species of the second one is replaced by a copy of the first one. Similarly, HGT gene-sweep events occur at each time step with a rate *p*_*h*_ (per patch, per time step). In such events, two random patches are selected. If the species of the first one carries the beneficial gene and the second one does not, then the gene is horizontally transferred and spreads into the second patch, without any displacement of species.

This model configuration corresponds to an evolutionary regime where the mantainance of diversity is slow compared to the time scale of fixation dynamics. For this regime we find that the number of populations (patches) carrying the beneficial gene follows a logistic growth (see Methods, Eq. 11) and after a time 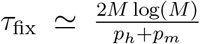 steps 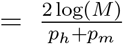 gen the diversity reaches an absorbing state where all the species in the metacommunity carry the advantageous gene. Mathematically, this evolutionary regime is defined by the condition *τ*_*fix*_ ≪ *τ*_*eq*_. The key aspect is the residual value of the diversity after the fixation of the beneficial gene.

Fig. 3B shows an example of the typical dynamics of the diversity after the introduction of the beneficial gene. The initial value of the diversity (*S*_*i*_ ≃ ⟨*S*_0_⟩) decreases after the introduction of the beneficial gene and reaches a new stationary value *S*_*f*_. We quantify the effect of the fixation of the beneficial gene by the “sweep parameter” 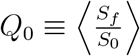. A value of *Q*_0_ ≃ 0 corresponds to a scenario of a genome-wide sweep across the metacommunity, while for *Q*_0_ *>* 0 some diversity is preserved.

We have derived an approximate analytical solution for the model dynamics in this regime

(i.e., when *τ*_*fix*_ ≪ *τ*_*eq*_, see methods), which leads to the following expression for the sweep parameter,

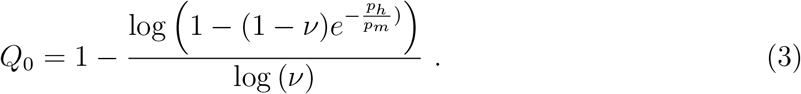

Eq. (3) is in good agreement with numerical simulations of the model (Fig. 3C,D). These results show that a full genome-sweep dynamics (*Q*_0_ = 0) can only be reached when *p*_*h*_*/p*_*m*_ → 0, i.e., when the HGT rate is completely negligible and the “invasion” dynamics of the species with the beneficial gene is the only relevant one. However, as soon as the HGT rate is non-negligible (for any positive value *p*_*h*_*/p*_*m*_ *>* 0), there is more than a single species within the metacommunity after the fixation of the beneficial gene. More specifically, for values of *p*_*h*_*/p*_*m*_ ≃ 0.1 and *ν* = 0.01, we already obtain *Q*_0_ ≃ 0.5, which means that the HGT-sweep rate is ten times slower than the typical migration-sweep time is sufficient to preserve (in the worst-case scenario) half of the diversity within a metacommunity.

### Gene-sweep dynamics under competing time scales

Having quantified how a metacommunity may preserve diversity under a gene sweep in the absence of diversity-restoring mechanisms (i.e., when *τ*_*fix*_ ≪ *τ*_*eq*_), we now study how diversity and multiple rounds of gene sweep may interact over longer time scales. Specifically, we consider a regime where the emergence and fixation dynamics of beneficial genes takes place on a time scale that is comparable with the relaxation time of the diversity-restoring mechanism, which occurs when *τ*_*fix*_ ⪆ *τ*_*eq*_ and is realized in our case as a neutral model (Fig. 4A). In this regime, three different evolutionary forces are acting at each time step: (i) innovation events, taking place at a rate *ν* (per patch, per time step), (ii) migration of a species with the beneficial gene into a patch that did not carry the gene (rate (1 − *ν*)*p*_*m*_, per patch, per time step), and (iii) transfer of a beneficial gene via HGT and sweep (rate (1 − *ν*)*p*_*h*_ per patch, per time step). In order to fully specify the model, we need to state how likely new species carry a beneficial gene. We assume that in an innovation event, the new species carries the beneficial gene with a probability *f*_0_(*t*) = *D*_*s*_(*t*)*/M*, i.e., equal to the fraction of populations (patches) carrying the beneficial gene at the time of the event. This assumption is justified when innovation represent migration events from a parallel metacommunity where the beneficial gene has the same frequency across patches.Finally, a species can migrate and sweep to another patch with its same genetic content (both species with the beneficial gene, or both without the gene) with a neutral rate 1 − *ν* (per patch, per time step).

**Figure 4.**
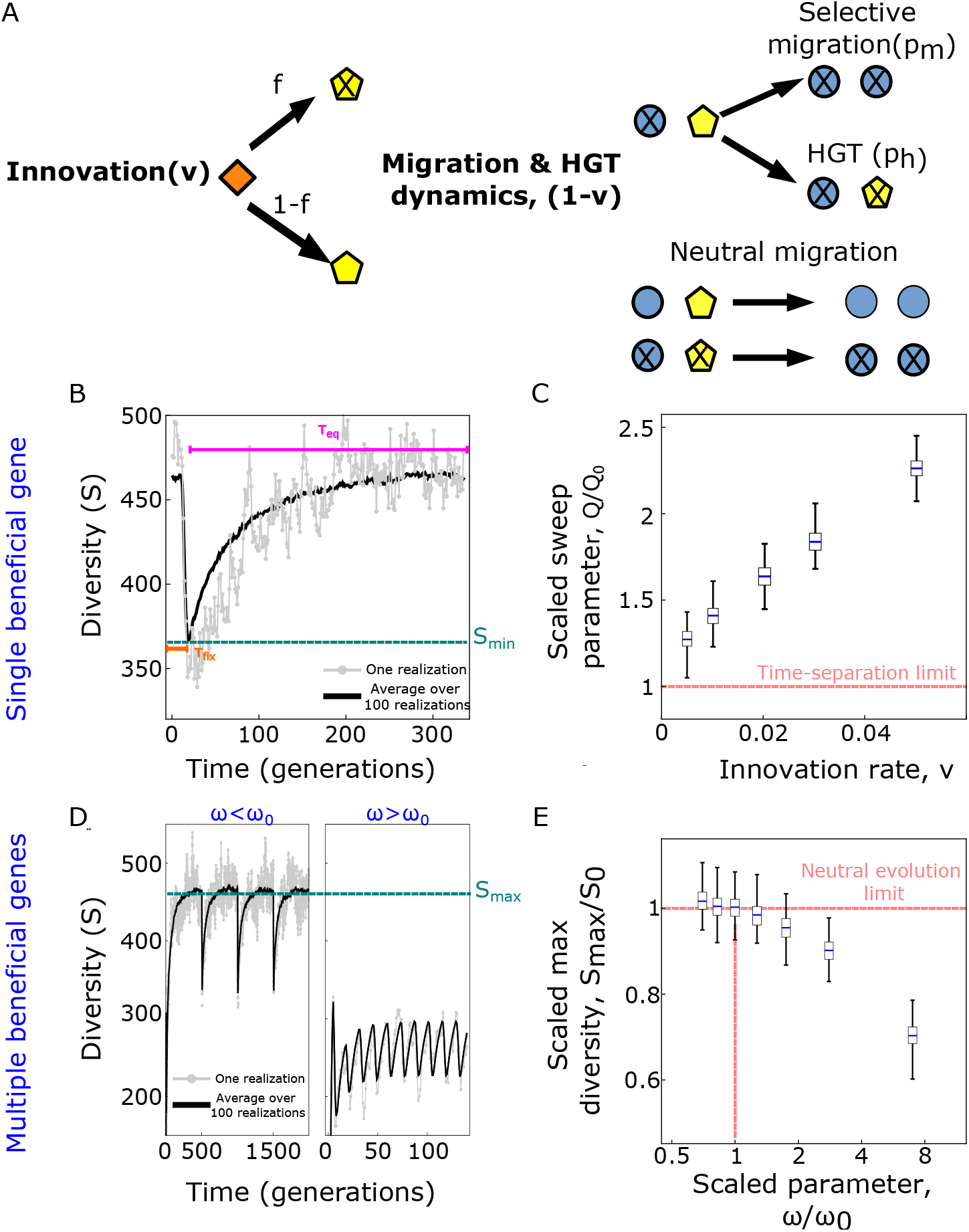
Competition of gene-sweep and diversity-restoring dynamics affects the minimal and maximal observed diversity. **A**. Spread of a beneficial gene over a metacommunity can compete with a diversity-restoring mechanism occurring with a rate *ν*. HGT-sweep and migration-sweep events take place with a joint rate 1 − *ν* and are realized by picking two random patches within the metacommunity. In innovations, new species will carry a beneficial gene with probability *f*_0_ equal to the fraction of populations (patches) within the metacommunity carrying the beneficial gene. Neutral migration-sweep events occur if both populations carry the beneficial gene, or neither of them does. If the first of the two populations carries the beneficial gene and the second does not, a (selective) migration-sweep event occurs with probability *p*_*m*_ (and a total rate (1 − *ν*)*p*_*m*_) while an HGT-sweep event occurs with probability *p*_*h*_ (and a total rate (1 − *ν*)*p*_*h*_), see Fig 3. **B**. The dynamics of diversity after the emergence of a beneficial gene is characterized by two time scales: (i) the time until the beneficial gene reaches fixation (*τ*_*fix*_), during which the diversity drops and (ii) an equilibration time (*τ*_*eq*_), restoring its initial value. **C**. Minimum diversity quantified by the scaled sweep parameter *Q/Q*_0_, where *Q* = ⟨*S*_*min*_*/S*_0_⟩, and *Q*_0_ is the sweep parameter (Fig. 3). The distribution of *Q/Q*_0_ shown as a box plot shows an increasing trend with increasing innovation rate *ν*. **D**. If multiple beneficial genes emerge periodically after a time 1/*ω*, diversity shows an oscillatory dynamics.**E**. The maximum diversity (*S*_*max*_, shown as a box plot) divided by the expected value under neutral biodiversity (*S*_0_), decreases after a critical value of the scaled parameter *ω/ω*_0_, where 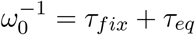 = *τ*_*fix*_ + *τ*_*eq*_). Other model parameters: *M* = 10000, *p*_*h*_ = 0.1, *p*_*m*_ = 0.9. All rates are per patch, per time step unless otherwise specified.

Fig.4B shows the typical dynamics of diversity after the introduction of a single beneficial gene. Diversity drops to a minimum value (*S*_*min*_), in a time-scale 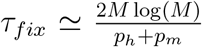steps = 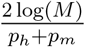 gen, for *ν* ≪ 1, similar to the case of the time-separation limit. However, in this case, the innovation dynamics restores the initial value of the diversity, on a time scale *τ*_*eq*_ ≃ *M/*(*ν*) steps = 1*/ν* gen. Moreover, because of the generation of new species during the fixation dynamics, the minimum value of the diversity is typically higher than the one observed in the time-separation limit, that corresponds to the mathematical limit *ν* → 0 and occurs when *tau*_*fix*_ ≪ *τ*_*eq*_ (cfr. Fig. 3). We have quantified this effect with numerical simulations of the model, using the sweep rate, now defined as

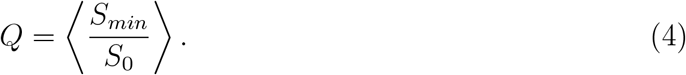

Fig. 4C shows that for innovation *ν* = 0 one obtains the same minimum for the diversity as in the limit case without any diversity-restoring mechanism, *Q* = *Q*_0_. Conversely, the minimal diversity is always higher than in absence of a diversity-restoring mechanisms (*Q > Q*_0_) under a positive innovation rate *ν >* 0. These observations arise due to the competition between the time scale of the diversity drop via the sweeping gene and the time scale of the diversity-restoring mechanism. If the restoring mechanism is sufficiently fast, the diversity-drop mechanism does not have time to reach its natural minimum value (due to the complete sweep) observed in Fig. 3B.

Due to the same competition of time scales, multiple acquisitions of beneficial genes may have consequences on the *maximal* diversity observed in the system. The mechanism is illustrated by Fig. 4DE. We have considered a regime where multiple beneficial genes arrive in the metacommunity with a constant frequency *ω* (per time step). Additionally, we assumed a fitness scenario where the last-emerged gene always carries the highest beneficial effect. In a simple simulation where beneficial genes arrive periodically after a time 1/*ω* the resulting dynamics of the diversity shows an oscillatory pattern due to successive gene-sweeps (Fig. 4D), with oscillations corresponding to rounds of gene-sweep and diversity-restoring dynamics. For beneficial genes arriving stochastically at a constant rate *ω*, the oscillations disappear, but the average diversity display similar behaviour (cfr. Fig.S1). If the rate of arrival of new beneficial genes is too high, the diversity-restoring mechanism does not have enough time to achieve its steady-state diversity. This process is illustrated by Fig. 4E, which quantifies the maximal diversity as a function of the arrival rate of new beneficial genes. In case *ω*^*−*1^ *> τ*_*fix*_ + *τ*_*eq*_, the time scale for the emergence of the beneficial rate is faster than the equilibration dynamics, hence the diversity cannot be restored to its initial value. Table S1) recapitulates the four different regimes for the qualitative behavior of the diversity in terms of two dimensionless quantities, *ω*(*τ*_*fix*_ + *τ*_*eq*_), and *τ*_*fix*_*/τ*_*eq*_. The first quantity compares the typical time between arrivals of the beneficial genes with the total time needed to fix them and to equilibrate the system by its neutral dynamics. The second is the ratio of the fixation time of the beneficial gene with the neutral equilibration time (which sets the dominant dynamics).

## DISCUSSION AND CONCLUSIONS

We have shown that an eco-evolutionary model given a metacommunity structure can support a “gene sweep” dynamics, without eliminating genome diversity. Instead, gene sweeps can lead to a moderate reduction of diversity even in the absence of diversity-restoring mechanism. Inclusion of a diversity-restoring mechanism (e.g, a neutral biodiversity model in our specific case) can increase the minimal observed diversity as well as the maximal one in the case of multiple sweeping beneficial genes. The mechanism by which a metacommunity maintains diversity under a gene sweep is compatible with small HGT rates compared to typical migration time scales. Unlike prior work, our model does not explicitly require additional ingredients such as frequency-dependent selection at the individual level, induced by genome-level processes or by ecological interactions.

In particular, the model outcomes rely on simple time-scale competition arguments. From this standpoint, our hypothesis is related in spirit to the classic proposition put forward by G. A. Hutchinson [24] to justify the very high observed microbiological diversity in samples of ocean pythoplankton (which was at odds with the principle of competitive exclusion, according to which the survival of a single species within a population should be privileged). To reconcile diversity with competitive exclusion, Hutchinson argued that if the time scale at which the exclusion principle is enforced were comparable to the time scale over which environmental conditions change significantly, a state of equilibrium would never be reached, and therefore there would be no predominance of a single species.

Our focus on a metacommunity is complementary to the approach assumed by the previous study by Niehuis and coworkers [6], which focused on intra-patch diversity of a *single* population and the role of migration of non-carrier individuals, favoring diversity in moderate amounts. The same study also showed that such migration effects are enhanced in a small metacommunity, made of multiple patches. In our model, all populations are connected, and the within-population dynamics are assumed to be fast and result in a single winner. More precisely, we took the conservative assumptions that (i) migration of carriers (which reduces the diversity) is the dominant process and that (ii) the presence of the beneficial gene on a patch, whether it is carried by a species invading the patch by migration or if is acquired by HGT, is sufficient to guarantee a full sweep of the population. Phenomena akin to those described by Niehuis and colleagues would increase the prediction of the residual diversity in our model. Thus, in light of their study, we can consider our estimates as lower bounds for diversity.

Apart from these differences, the mechanisms described by our work are conceptually similar to the ones discussed by Niehius and coworkers, in that they are a result of the balance between HGT and migration rates. As noted in ref. [6], these mechanisms lead to an *effective* frequency-dependent selection (which in our case acts completely at the level of populations within a metacommunity, not on individuals). However, we note that this dependency has a different origin than the processes hypothesized by Takeuchi and coworkers [14] (which act at the level of an individual within a population). In the scenario assumed by Takeuchi and colleagues, the diversity is favored by ubiquitous and diverse deleterious loci that are linked to the acquired beneficial gene. In such cases, the diversity of bacterial species should be capped by the number of deleterious linked effects (e.g. phage diversity). If these linked alleles can be quantified in data, they should be linked to residual diversity after a gene sweep.

Importantly, even though they are conceptually different, the two hypotheses are not mutually exclusive, and possibly can both be detected in data or addressed in controlled experiments. To address negative frequency-dependent selection due to linkage, genomic analysis of microbial communities undergoing gene sweeps should be able to isolate the linked deleterious loci that co-occur with the beneficial gene in each species or strain. In order to test the role of a patchy community in restoring diversity, controlled experiments could induce gene sweeps in a laboratory metacommunity with varying densities of patches. Experimental systems that might allow this are conceivable today [18], although complex spatio-temporal processes might complicate considerably the experimental scenario compared to the simple, purely conceptual, model proposed here [25, 26]. Despite these limitations, future studies of genomic data may be able to differentiate a gene-specific sweep with or without high HGT based on an analysis of additional selective forces in other portions of the genome.

## METHODS

### Within-patch and between-patches fixation dynamics

Let us first consider an individual patch supporting the growth of *N* clonal individual. Let us fix the timescales so that the generation rate (time) equals to 1. We consider a mutation with selective advantage *s* and assume 1*/N* ≪ *s* ≪ 1. The fixation probability of such a mutation equals 1 − exp(−*s*) ≈ *s*, and the typical fixation timescales (intra-patch sweep time) equals *τ*_*patch*_ ∼ 1*/s* log(*Ns*).

Successful migrations and HGT of a beneficial genome (gene in the case of HGT) with selective advantage *s* occurs with rates *p*_*m*_ and *p*_*h*_ respectively. Assuming that the timescales of intra- and interpatch dynamics are separated, these rates can be decomposed into two contributions: a basal migration / HGT rates, equal to *µ*_*m*_ and *µ*_*h*_ respectively, and the fixation probability equal to *s*. Therefore we have *p*_*m*_ = *sµ*_*m*_ and *p*_*h*_ = *sµ*_*h*_.

The other processes shaping the communities are the innovations and the displacement of a species by another one from a patch due to neutral migration. We define *µ*_*I*_ as the rate of arrival of a new species in the patch. Assuming neutrality, the probability of fixation in a patch is 1*/N*. We have therefore that a new species is introduced with rate *µ*_*I*_*/N* and the displacement of a species on a patch via migration occurs with rate *µ*_*m*_*/N*.

In all our derivations we assume a time-scale separation between these three processes: fixation of beneficial mutations in a single patch (with rate ≈ *s/* log(*Ns*)), migration/HGT across patches (with rates *sµ*_*m*_ and *sµ*_*h*_), and neutral processes and diversity innovations (rates *µ*_*m*_*/N* and *µ*_*I*_*/N*):

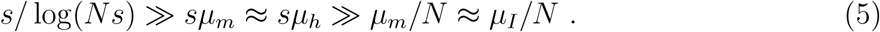

Together with the assumption 1*/N* ≪ *s* ≪ 1, this condition impose the constraints

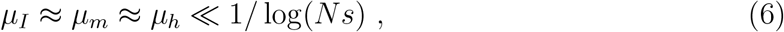

where time is measured in generations (here defined as the typical doubling time of individuals within a patch).

### Expected diversity for the neutral model

This section focuses on a model configuration where only two evolutionary forces are present: (i) innovation events, taking place at rate *ν* (per patch, per time step) and (ii) neutral migrations (rate 1 − *ν*, per patch, per time step). This configuration corresponds to the standard Hubbell’s model [20]. In the following, we will show that, using analytical results known for such model classes [20, 27], one can easily compute the expected value of the diversity at equilibrium, in the neutral scenario (Eq.1).

Hubbel’s model can be mapped into a urn process, where each patch is represented by a ball with a color corresponding to its species, and it can be shown [20, 27] that the distribution of the species abundance at equilibrium is given by Ewens’s sampling formula, for the distribution of alleles under neutral mutations [28], in the context of population genetics

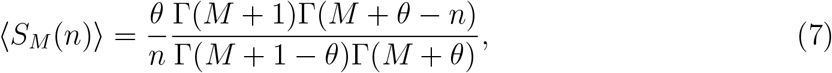

where 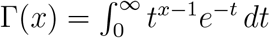 is the standard Gamma function.

This distribution defines the expected number of species occupying exactly *n* patches, and is specified in terms of the model parameter 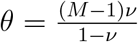, called the fundamental biodiversity number. The expected number of species co-existing in the metacommunity at equilibrium is then obtained by summing over all the elements of this distribution [27]

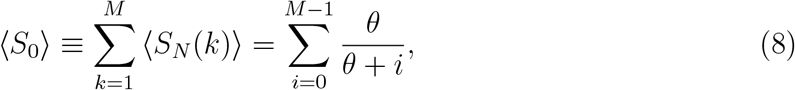

In the limit *M* ≫ 1, we can replace the sum with an integral,

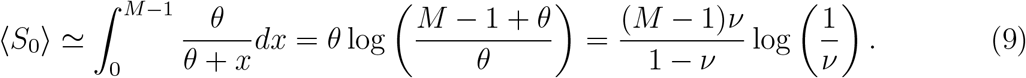

In the limit of small values of *ν* and by replacing *M* − 1 → *M*, we obtain the approximated solution of Eq.(1)

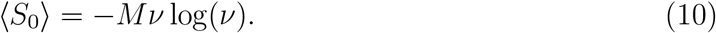

### Model dynamics during the spread of a beneficial gene, without diversity-maintenance mechanism

In this section, we focus on the model variant described in Fig.2, where a beneficial gene is introduced in the metacommunity and can spread through to invasions and HGT events, with a total rate *p*_*h*_ + *p*_*m*_ (per patch, per time step).

First, we focus on the the time-evolution of the average number of patches with populations carrying the beneficial genes (*B*(*t*)), which is described by a logistic growth

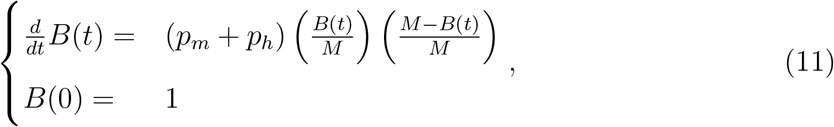

where time is measured in time steps from the introduction of the beneficial gene, in the continuous limit approximation. The expected time to the fixation of the beneficial gene in the metacommunity (*τ*_*fix*_) can be computed using the solution of Eq.11, which reads

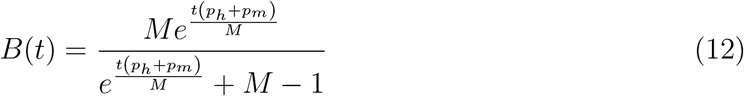

and imposing the condition *B*(*τ*_*fix*_) = *M* − 1, which is fulfilled by

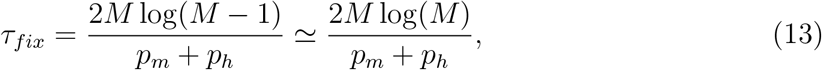

where time is counted in time steps.

Next, we focus on the extinction dynamics of a species without the beneficial gene. To compute the extinction probability of such species, we start by considering the dynamics of the number of patches without the beneficial gene and with species *s* (*D*_*s*_(*t*)). In the model configurations considered in this context, this class of species can only decrease over time because of the invasion of species carrying the beneficial gene. The time evolution of *D*_*s*_(*t*) and given by

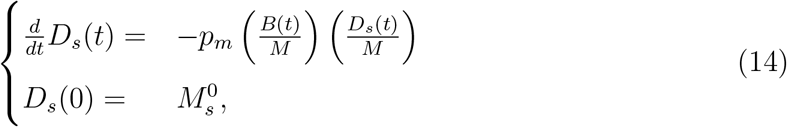

since invasion events occur with a rate *p*_*m*_, and cause a reduction of *D*_*s*_(*t*) if one of the two populations picked randomly belongs to *s* (i.e., chosen with probability *D*_*s*_(*t*)*/M*) and the other one carries the beneficial gene (i.e.,chosen with probability *B*(*t*)*/M*). Here, time is measured in time steps from the introduction of the beneficial gene, in a continuous limit approximation. The solution to Eq.(14) reads

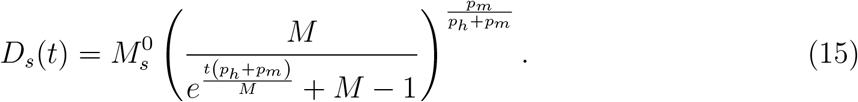

Similarly, the probability that, at time *t*, one of the species *s* acquires the beneficial gene by HGT is given by the product of the HGT rate, *p*_*h*_ and the probability that one of the two populations picked randomly belongs to *s* (i.e., chosen with probability *D*_*s*_(*t*)*/M*) and the other one carries the beneficial gene (i.e.,chosen with probability *B*(*t*)*/M*)

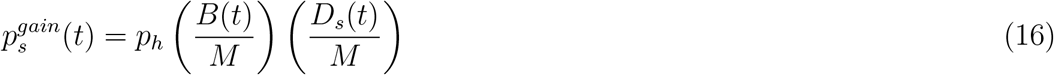

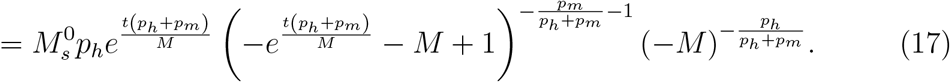

The expected number of species *s* that, by time *t*, have gained the beneficial gene is

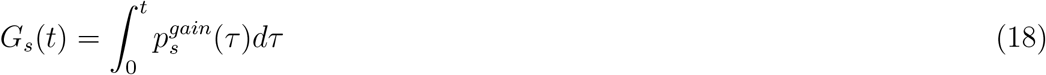

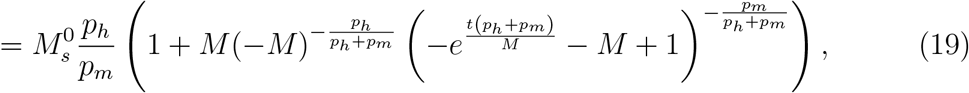

and the expected number of species *s* that have gained the beneficial gene at any point in time can be computed as

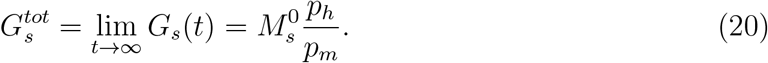

The probability of extinction for species *s* can be computed assuming a Poisson distribution for the number of species *s* that have gained the beneficial gene. This distribution has mean value 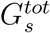, Hence, the probability to have gained 0 genes (i.e., to became extinct) reads

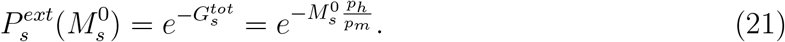

Finally,the sweep parameter, i.e., expected reduction of the biodiversity after the fixation of the beneficial gene, is obtained by averaging 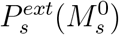 over the probability distribution of the number of patches populated by the same species (*P* (*M* ^0^))

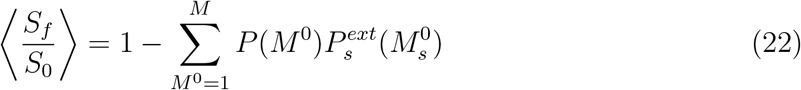

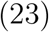

The probability distribution *P* (*M* ^0^) is the normalized version of Eq.(7), and, in the limit of large metacommunity *M* ≫ 1 and small innovation rate *ν* ≪ 1, is well approximated by the Fisher log series[27, 29]

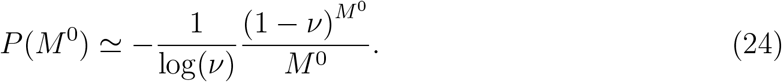

Under these assumptions, the sweep parameter can be computed analytically and reads

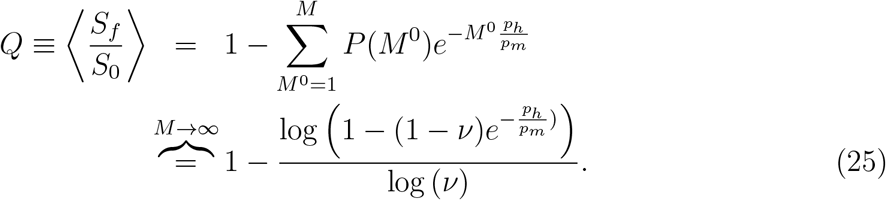

## Code Availability

The code used for the simulations is provided in the GitHub repository: https://github.com/eddbell/MetapopulationModel

## Aknowledgements

We would like to thank Kunihiko Kaneko for useful discussions.

---

**Table S1.** Expected values of diversity in all model regimes. This table summarizes the different regimes of the expected value of the biodiversity as a function of two dimensionless variables *ω*(*τ*_*fix*_ + *τ*_*eq*_), and *τ*_*fix*_*/τ*_*eq*_, including the three main time scales of the model: (i) the equilibration time of the system (*τ*_*eq*_), (ii) the fixation time of the beneficial gene (*τ*_*fix*_) and (iii) the time between arrival of beneficial genes 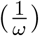. In each regime we illustrate the (i) expected maximum and minimum biodiversity (*S*_max_ and *S*_min_, expressed in terms of the expected neutral value of the diversity *S*_0_ (Eq.1) and of the sweep parameter *Q*_0_ (Eq.3) and (ii) the Figure showing the corresponding numerical results.

| Regime | Conditions on time scales | Expected diversity | Figure of reference |
| --- | --- | --- | --- |
| Neutral evolution | / | $\langle S \rangle = \langle S_0 \rangle$ | Fig.2 |
| Time separation<br>(No diversity-maintenance mechanism) | $\tau_{fix}/\tau_{eq} \ll 1$ | $S_{max} \geq S \geq S_{min}$<br>$\langle S_{max} \rangle = \langle S_0 \rangle$<br>$\langle S_{min} \rangle = Q_0 \langle S_0 \rangle$ | Fig.3 |
| Time scale competition | $\tau_{fix}/\tau_{eq} \gtrsim 1$ and $\omega(\tau_{fix} + \tau_{eq}) < 1$ | $S_{max} \geq S \geq S_{min}$<br>$\langle S_{max} \rangle = \langle S_0 \rangle$<br>$\langle S_{min} \rangle > Q_0 \langle S_0 \rangle$ | Fig.4 B-C |
| | $\tau_{fix}/\tau_{eq} \gtrsim 1$ and $\omega(\tau_{fix} + \tau_{eq}) > 1$ | $S_{max} \geq S \geq S_{min}$<br>$\langle S_{max} \rangle < \langle S_0 \rangle$<br>$\langle S_{min} \rangle > Q_0 \langle S_0 \rangle$ | Fig.4 C-D |

**Figure S1.**
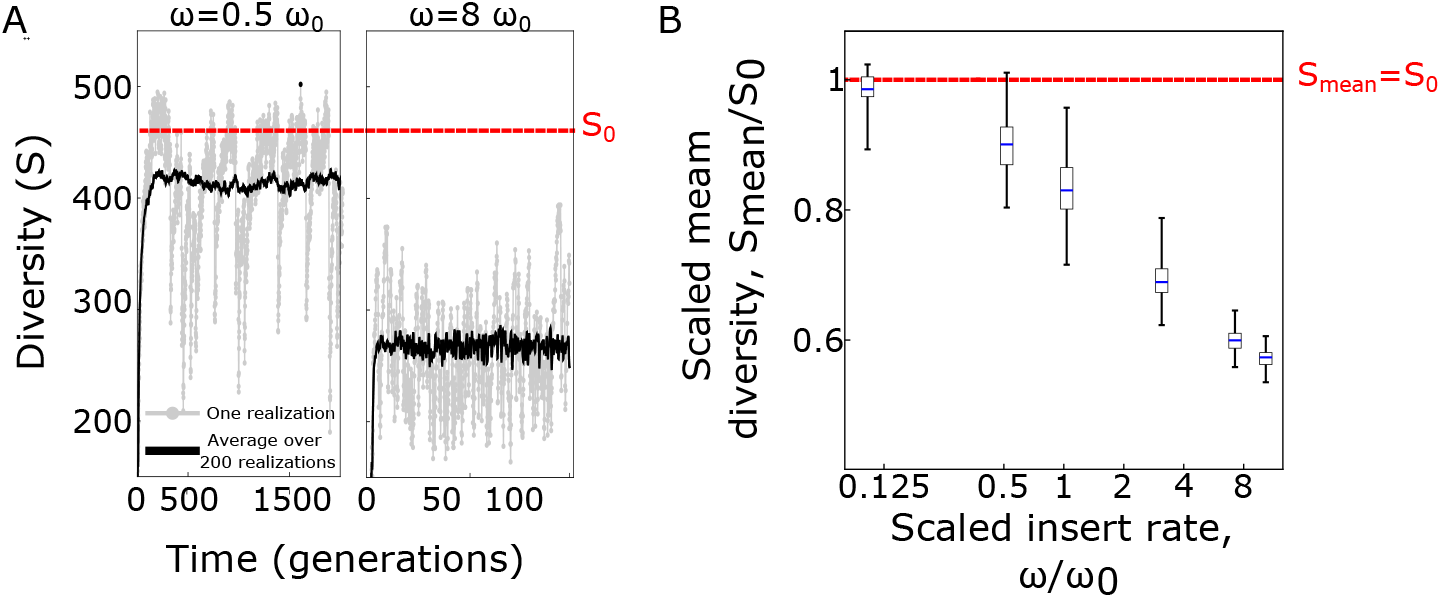
Dynamics of the average diversity when beneficial genes emerge at a constant rate. **A**. We show here two examples of simulations use to investigate the dynamics of the diversity in the same model regime as Fig.4, when beneficial genes emerge stochastically at a constant rate *ω* (values of *ω* specified above each plot, in terms of 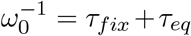 = *τ*_*fix*_ + *τ*_*eq*_)). In this case no oscillations are observed, and the dynamics of the diversity is captured by its average value *S*_*mean*_.**B**. The average diversity (*S*_*mean*_, shown as a box plot) divided by the expected value under neutral biodiversity (*S*_0_) shows a monotonic decrease with increasing value of *ω/ω*_0_, where 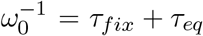). Other model parameters: *M* = 10000, *p*_*h*_ = 0.1, *p*_*m*_ = 0.9. All rates are per patch, per time step unless otherwise specified.

